# Genetically encoded nanostructures enable acoustic manipulation of engineered cells

**DOI:** 10.1101/691105

**Authors:** Di Wu, Diego Baresch, Colin Cook, Dina Malounda, David Maresca, Maria Paulene Abundo, David Reza Mittelstein, Mikhail G. Shapiro

## Abstract

The ability to mechanically manipulate and control the spatial arrangement of biological materials is a critical capability in biomedicine and synthetic biology. Ultrasound has the ability to manipulate objects with high spatial and temporal precision via acoustic radiation force, but has not been used to directly control biomolecules or genetically defined cells. Here, we show that gas vesicles (GVs), a unique class of genetically encoded gas-filled protein nanostructures, can be directly manipulated and patterned by ultrasound and enable acoustic control of genetically engineered GV-expressing cells. Due to their differential density and compressibility relative to water, GVs experience sufficient acoustic radiation force to allow these biomolecules to be moved with acoustic standing waves, as demonstrated within microfluidic devices. Engineered variants of GVs differing in their mechanical properties enable multiplexed actuation and act as sensors of acoustic pressure. Furthermore, when expressed inside genetically engineered bacterial cells, GVs enable these cells to be selectively manipulated with sound waves, allowing patterning, focal trapping and translation with acoustic fields. This work establishes the first genetically encoded nanomaterial compatible with acoustic manipulation, enabling molecular and cellular control in a broad range of contexts.

## INTRODUCTION

The ability to remotely manipulate and pattern cells and molecules would have many applications in biomedicine and synthetic biology, ranging from biofabrication^1^ and drug delivery^2^ to noninvasive control of cellular function^3–5^. Ultrasound offers unique advantages in such contexts over optical, magnetic and printing-based approaches due to its non-invasiveness, functionality in opaque media, and its relatively high spatial precision on the µm scale. Acoustic radiation force (ARF) allows ultrasound to manipulate objects whose density or compressibility differ from their surrounding medium. This capability has been used to manipulate, pattern and sort synthetic particles and large mammalian cells, for example by using acoustic standing waves to create stable attractors for such objects or separate them in microfluidic devices^6^. However, due to their small size and weak acoustic contrast relative to water, biomolecules have not been manipulated directly with ARF. Furthermore, the similarity in acoustic properties between cells has made it challenging to separate cells of a similar size based on their genotypes.

To address these limitations, we hypothesized that gas vesicles (GVs), a unique class of genetically encoded air-filled protein nanostructures, could experience ARF and enable the acoustic manipulation of GV-expressing cells. GVs are protein-shelled nanostructures with hydrodynamic diameters on the order of 250 nm (Fig. 1, a-b) which evolved in aquatic photosynthetic microbes as a means to achieve buoyancy for improved access to sunlight^7^. GVs consist of a physically stable hollow compartment enclosed by a 2 nm-thick protein shell that is permeable to gas but excludes liquid water. Based on their unique physical properties, GVs were recently developed as genetically encodable and engineerable contrast agents for non-invasive imaging^8–13^. However, the ability of GVs to serve as actuators of ARF has not been tested.

**Fig. 1.**
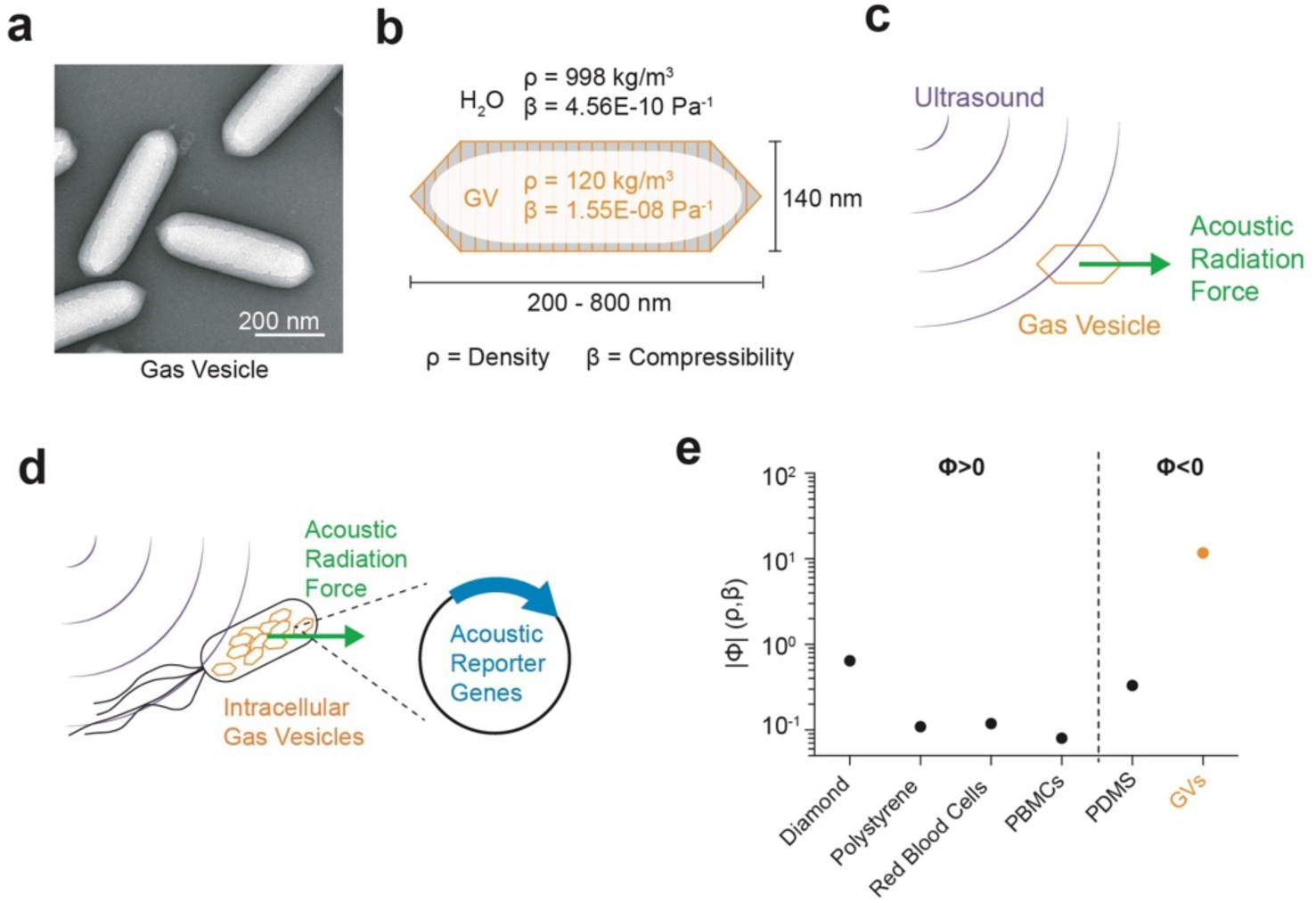
Gas vesicles as nanotransducers of acoustic radiation force. **a**, Transmission electron microscopy image of representative GVs from *Anabaena flos-aquae*. **b**, Schematic drawing of a GV, showing its effective density (ρ) and compressibility (β) relative to that of the surrounding water. **c**, Illustration of a GV experiencing acoustic radiation force due to applied ultrasound. **d**, Illustration of a genetically modified bacterium experiencing enhanced acoustic radiation force due to the expression of GVs inside the cell. **e**, Estimated magnitude of the acoustic contrast factor, |Φ|, of GVs and several common materials used in acoustic manipulation. Materials to the left and right of the vertical dashed line exhibit positive and negative acoustic contrast in water, respectively. PBMCs, peripheral blood mononuclear cell. PDMS, polydimethylsiloxane.

We hypothesized that GVs’ differential density and compressibility relative to aqueous media would allow these nanostructures to experience significant ARF (Fig. 1c), and that cells genetically engineered to express GVs would experience enhanced radiation force due to changes in their acoustic properties (Fig. 1d). We further hypothesized that the resulting forces would act in the opposite direction from other biomaterials, which are generally denser than water, allowing selective acoustic manipulation. In this study, we test these fundamental hypotheses and demonstrate the use of GVs in dynamic patterning, multiplexed acoustic manipulation, measurement of acoustic pressure and cellular actuation.

## RESULTS

### Gas vesicles experience direct acoustic radiation force

To estimate the expected ARF acting on GVs, we modeled them as approximately spherical particles with an effective density of 120 kg/m^3^ (ref. ^14^) and compressibility of 1.55E-8 Pa^−1^ (ref. ^15^). Because both of these values are radically different from water (Fig. 1b), we predicted that GVs would have a strongly negative acoustic contrast in aqueous media, with a contrast factor of –11.7 (Fig. 1e, Eq. 1 in Methods). This differs in both sign and magnitude from most materials used in acoustic manipulation. For example, polystyrene and red blood cells have acoustic contrast factors on the order of +0.1. The exceptionally large acoustic contrast of GVs suggested that, despite their small size, these biomolecules could be manipulated with ultrasound at typical ARF frequencies and energy densities (MHz and ~10-100 J/m^3^)^16^.

To test the ability of GV nanostructures to be manipulated with ARF, we purified GVs from the cyanobacterium *Anabaena flos-aquae* (Ana GVs), chemically labeled them with a fluorescent dye, and imaged them in suspension inside a microfluidic channel coupled to a bulk piezoelectric resonator operating at 3.8 MHz (Fig. 2a). The channel width of 200 μm represents a half-wavelength at this frequency, resulting in a pressure node at its center and antinodes (areas of highest pressure) at each wall (Fig. 2b). As expected based on their negative acoustic contrast, GVs readily migrated to the pressure anti-nodes upon ultrasound application (Fig. 2, c-d). As a control, we imaged GVs that were collapsed before the experiment with hydrostatic pressure (**Supplementary Fig. 1**). Neither collapsed GV nor similarly-sized polystyrene tracer nanoparticles, included as an additional control, migrated in the acoustic field.

**Fig. 2.**
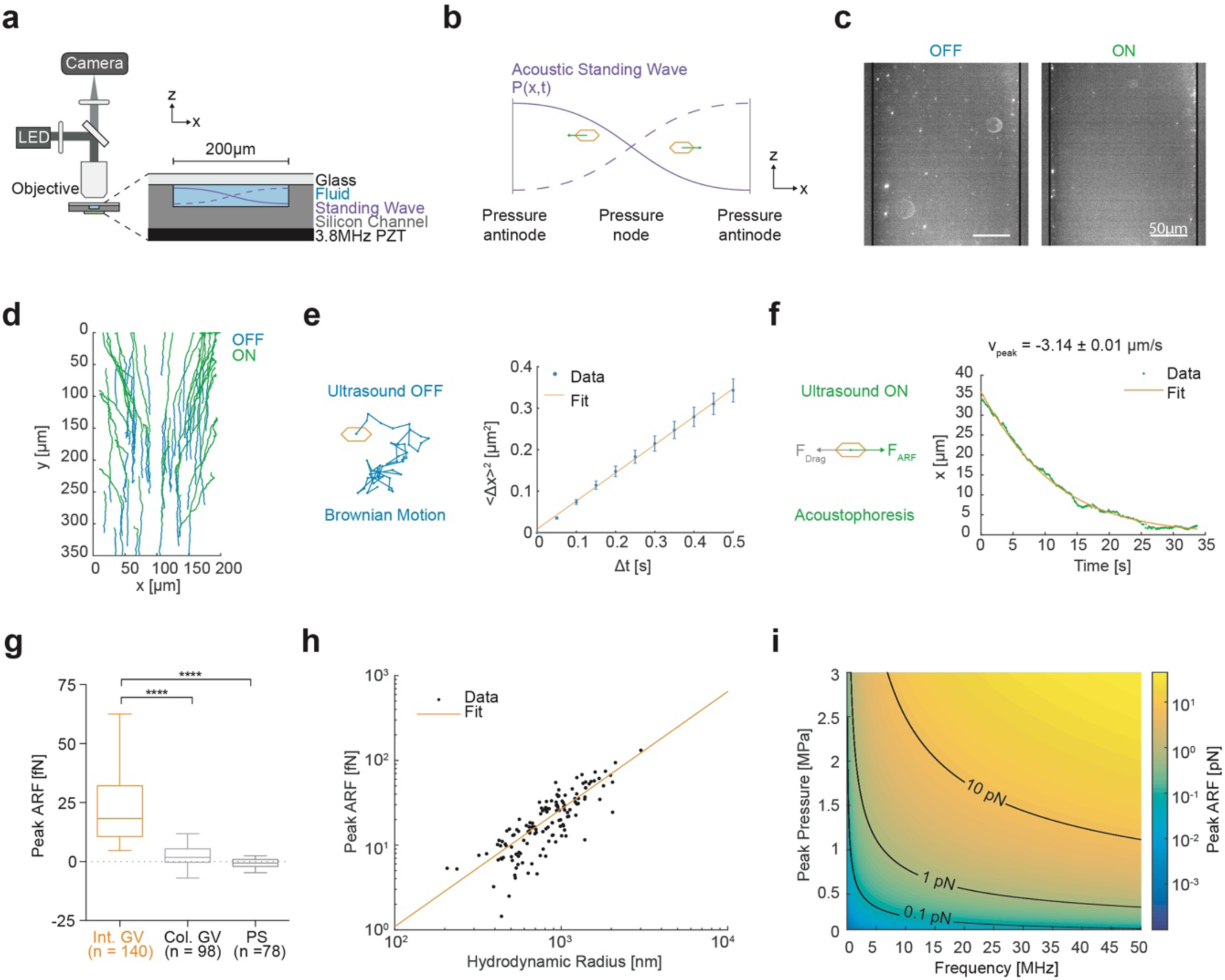
Gas vesicles experience direct acoustic radiation force. **a**, Diagram of the acoustic standing wave setup. A piezoelectric element is coupled to an etched silicon channel whose width is half the acoustic wavelength to generate a standing wave along the x-direction. The channel depth is 47 µm. Particles suspended in an aqueous solution are imaged using an epifluorescence microscope. LED, light-emitting diode. PZT, lead zirconate titanate. **b**, Illustration of the expected migration direction of GVs towards the pressure antinodes of an acoustic standing wave, due to their negative acoustic contrast. **c**, Fluorescence images of GVs inside the microfluidic channel before ultrasound (OFF) and 100 seconds after ultrasound has been turned on (ON). **d**, Representative single particle trajectories of GVs before (blue) and during (green) ultrasound application. **e**, Illustration of Brownian motion (left) and representative single-particle mean square displacement curve used to determine the diffusivity of the particle (right). **f**, Illustration of particle acoustophoresis (left) and representative single-particle trajectory in the *x* direction during ultrasound application, used to determine the peak particle velocity (right). **g**, Peak acoustic radiation force of intact GVs (24.5 ± 1.7 fN, n=140), pressure-collapsed GVs (2.0 ± 0.7 fN, N=98), and 200-nm polystyrene particles (−0.6 ± 0.4 fN, N=78). Box-and-whisker plots show the 5-95 percentile, the 25-75 percentile and the median of the distribution. Mann-Whitney test (****: p<0.0001). **h**, Peak ARF of GV particles as a function of hydrodynamic radius, fitted to a fractal clustering model (force-mobility exponent = 1.39±0.06; R^2^ = 0.744). **i**, Predicted ARF on a single GV across a range of acoustic parameters.

Next, we quantified the ARF acting on GV particles in solution using single-particle tracking (Fig. 2d). The Brownian motion of each particle before ultrasound application was used to determine its mobility and hydrodynamic size (Fig. 2e, Eq. 2 & 5 in Methods). For the same particle, its motion within the acoustic field during ultrasound application was fitted to an equation accounting for the spatial field profile (Eq. 4 in Methods), allowing us to determine the peak particle velocity (Fig. 2f). The maximum ARF acting on GV particles was then determined by a balance with hydrodynamic drag, and measured to be 24.5±1.7 fN under the acoustic parameters used in this measurement (Fig. 2g). In contrast, control particles showed no substantial ARF.

Non-specific association of individual GVs within the microfluidic channel resulted in tracked particles having a range of hydrodynamic radii larger than expected from a single GV. Therefore, to estimate the ARF acting on a single GV we plotted the dependence of the ARF on the hydrodynamic radius of the clusters and fitted it with a power law function accounting for fractal clustering^17^ (Fig. 2h, Eq. 6 in Methods, force-mobility exponent = 1.39±0.06; R^2^ = 0.744). Given the acoustic energy applied in this experiment (0.25±0.02 J/m^3^, **Supplementary Fig. 2**), this single-particle force corresponds to an acoustic contrast factor of –15±9, consistent with our theoretical estimate of –11.7 (Fig. 1e). Using this contrast factor, we can predict the ARF on a single GV across a range of typical acoustic parameters^16^ (Fig. 2i), with the expected force spanning from 0.01 to 10 pN. Forces of this magnitude are more than sufficient to overcome Brownian motion, as shown in our experiments, and are relevant to many biomolecular and cellular interactions^18^. Overall, these results establish the fundamental ability of GVs to be manipulated with acoustic fields.

### Gas vesicle collapse allows multiplexed manipulation and in situ pressure measurement

Having established the ability of GVs to experience strong ARF, we sought to take advantage of another unique property of these nanostructures: their ability to be collapsed at specific acoustic pressures determined by genetic engineering (Fig. 3, a-b)^8,13^. We hypothesized that this property would allow GVs to serve as probes for *in situ* pressure measurement, and that multiple GV types with different critical collapse pressures could be differentially manipulated in space.

**Fig. 3.**
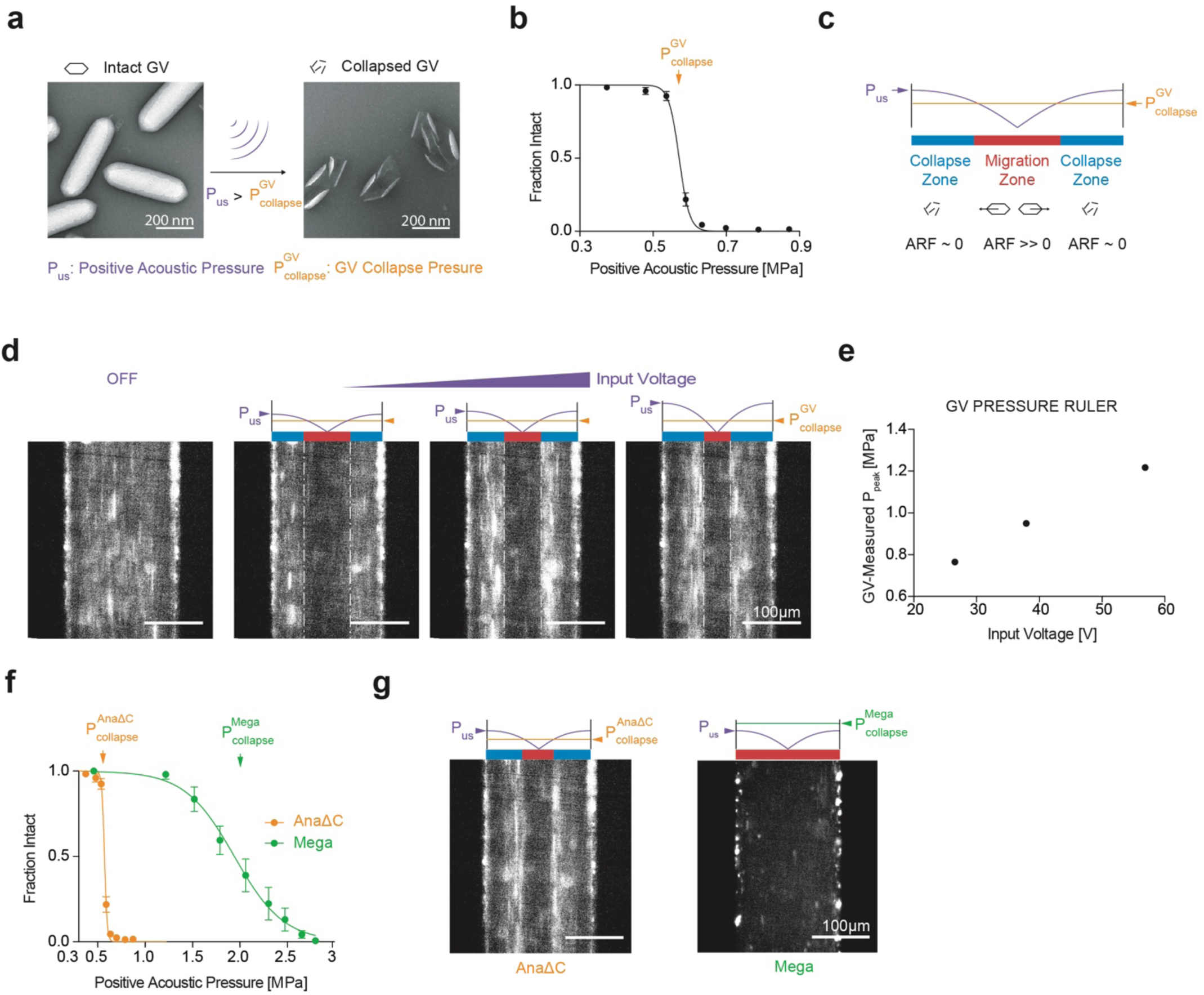
GV collapse enables *in situ* pressure sensing and multiplexed acoustic manipulation. **a**, TEM images of intact and collapsed Ana GVs. Collapse occurs when the positive acoustic pressure exceeds the critical collapse pressure of the GV. **b**, Acoustic collapse profile of AnaΔC GVs. The critical collapse pressure is determined to be the pressure at which 50% of the GVs have been collapsed. Data adapted from ref.^13^. **c**, Illustration of the expected behavior of GVs inside a microfluidic channel with a half-wavelength standing wave. GVs in regions with acoustic pressures lower than their critical collapse pressure migrate towards regions of higher pressure due to ARF, while GVs in regions with pressure above their critical threshold collapse and therefore remain stationary. The boundary between laterally migrating and stationary GVs indicates a pressure corresponding to the GVs’ critical collapse pressure. P_US_ indicates the temporal peak pressure. **d**, Fluorescence images of GVs inside a microfluidic channel in the presence of an acoustic field driven with increasing voltage. **e**, Maximal pressure in the acoustic device as a function of input voltage, determined using the images in (d). **f**, Acoustic collapse pressure curves of AnaΔC and Mega GVs. Data adapted from refs.^13,36^. **g**, Fluorescence images of either AnaΔC or Mega GV solutions experiencing the same acoustic field, with the peak driving pressure of 1.2 MPa selected to be above the critical collapse pressure of AnaΔC GVs, but below that of Mega GVs.

The measurement of acoustic pressure inside enclosed microstructures such as microfluidic channels is a major challenge in the field of acoustofluidics. Conventional methods to indirectly calibrate devices by inferring pressure profiles from the ARF exerted on single-particle standards are often highly laborious^19,20^. In contrast, we hypothesized that the pressure inside a microfluidic channel could be easily determined by visualizing the collapse location of GVs within the channel. In this approach, GVs in regions with acoustic pressures lower than their critical collapse pressure would migrate towards regions of higher pressure due to ARF, while GVs in regions with pressure above their critical threshold would collapse and therefore remain stationary. The boundary between migrating and stationary GVs corresponds to their collapse pressure (Fig. 3c). Since the relative pressure across the channel follows a known sinusoidal function, a single image revealing the location of GV collapse provides the entire standing wave pressure profile inside the channel.

To test this possibility, we imaged an engineered variant of Ana GVs (AnaΔC), whose acoustic collapse pressure (Fig. 3b) has been engineered to be lower than wild-type Ana GVs by removing the outer scaffolding protein GvpC^13^. We applied three different driving voltages to the piezoelectric element coupled to our microfluidic channel and imaged the steady-state distribution of GVs inside the channel. The expected pattern of nanoparticle discontinuity was observable starting with the lowest applied voltage (Fig. 3d). As we increased the driving voltage, the location of this discontinuity shifted inwards, consistent with the expected increase in acoustic pressure (Fig. 3d and **Supplementary Movie 1**). Using the locations of this discontinuity, we were able to calibrate the peak acoustic pressure in the device as a function of the driving voltage (Fig. 3e).

In addition to allowing GVs to serve as *in situ* pressure rulers, we hypothesized that spatially distinct collapse points would allow GVs with different characteristic collapse pressures to be differentially manipulated at the microscale. This is often desirable for example to enable separate visualization or multiplexed separation of analytes. To test this possibility, we imaged either AnaΔC GVs or heterologously expressed *B. megaterium* GVs (Mega GVs), which have critical collapse pressures of 0.6 MPa and 1.9 MPa, respectively (Fig. 3f). These GVs were subjected to a standing wave with a maximum acoustic pressure of 1.2 MPa, which should collapse AnaΔC but not Mega GVs. As expected, we observed that the two GV populations followed distinct migration patterns inside the acoustic field (Fig. 3g). These results establish the unique acoustofluidic capabilities provided by GVs’ engineerable acoustic collapse behavior.

### Heterologous expression of GVs enables selective acoustic manipulation of living cells

After establishing purified GVs as a biomolecular material for acoustic manipulation, we tested the ability of these genetically encodable nanostructures to act as a driver of ARF response in living cells. This possibility is based on the fact that GV expression significantly reduces the average density of the cell, resulting, for example, in the floatation of GV-expressing bacteria in water^12^. In combination with an anticipated increase in average cellular compressibility, this is expected to change the acoustic contrast of the cells from +0.07 to –1.1, flipping the sign of their acoustic contrast from positive to negative and increasing its magnitude by approximately 15-fold.

We tested this hypothesis by heterologously expressing intracellular GVs in *E. coli* using a recently developed genetic construct, *arg1*, comprising of a combination of 13 genes from *A. flos-aquae* and *B. megaterium* (Fig. 4, a-b).^12^ After enriching for high expression using centrifugation, the cells were labeled with a fluorescent dye to enable live cell tracking. *arg1*-expressing cells or control cells with pressure-collapsed intracellular GVs were then subjected to acoustic standing waves using the microfluidic device depicted in Fig. 2a. Remarkably, while control cells showed no response to the applied acoustic field, the genetically modified *arg1*-expressing cells containing intact intracellular GVs quickly migrated to pressure antinodes at the channel wall (Fig. 4c and **Supplementary Movie 2**). This result confirms that GV expression results in cells having a negative contrast factor, which is opposite from normal cells (Fig. 1e), and shows that the magnitude of this contrast factor is substantially larger than for wildtype controls, since under the same acoustic conditions the control cells did not migrate to the pressure node.

**Fig. 4.**
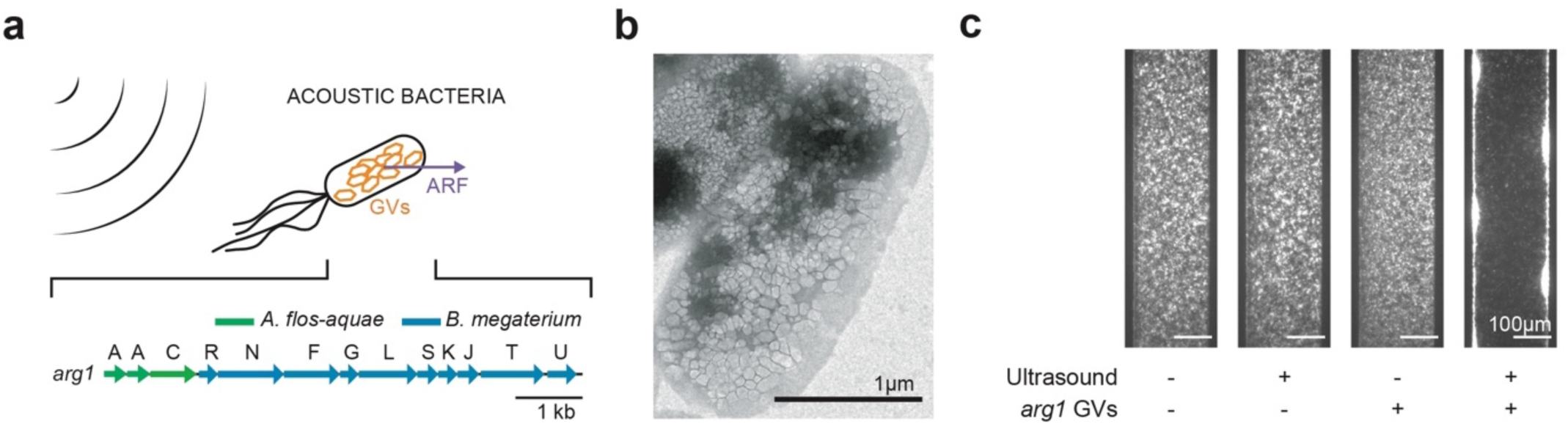
Gas vesicle expression in living cells enhances and changes the sign of their response to ARF. **a**, Schematic drawing of genetically modified *E.coli* experiencing an enhanced ARF due to the expression of intracellular GVs as acoustic reporter genes. **b**, TEM image of *E.coli* containing intracellular GVs upon expression of *arg1*. **c**, Fluorescence images of *E.coli* inside the microfluidic channel with either intact (+) or collapsed (−) intracellular GVs, either in the presence or absence of applied ultrasound.

Having established that GV-expressing cells experience strong ARF towards areas of high acoustic pressure, we asked whether this capability would enable the trapping and spatial patterning of living cells. Considerable interest exists in the use of engineered cells as patterned components of living materials for biomedical uses such as tissue engineering and as self-healing and actively reconfigurable materials in non-biomedical applications^21,22^. However, few methods exist to dynamically configure the location of cells in 3-D space. In contrast, ARF in the form of engineered standing and traveling waves has been used to create complex 2-D and 3-D arrangements^23–25^.

We hypothesized that ARF combined with GV expression would allow engineered cells to be patterned in a precise and rapid manner. To test this basic concept, we generated a standing wave pattern of repeating pressure antinodes in a specially designed acoustic chamber by using an unfocused 5 MHz transducer reflected by glass (Fig. 5a). Imaging the cells using fluorescence microscopy, we observed that engineered cells readily adopted the desired pattern in solution, and that changing the ultrasound frequency allows the spatial pattern of these cells to be dynamically reconfigured on the timescale of seconds (Fig. 5b, **Supplementary Fig. 3** and **Supplementary Movie 3**).

**Fig. 5.**
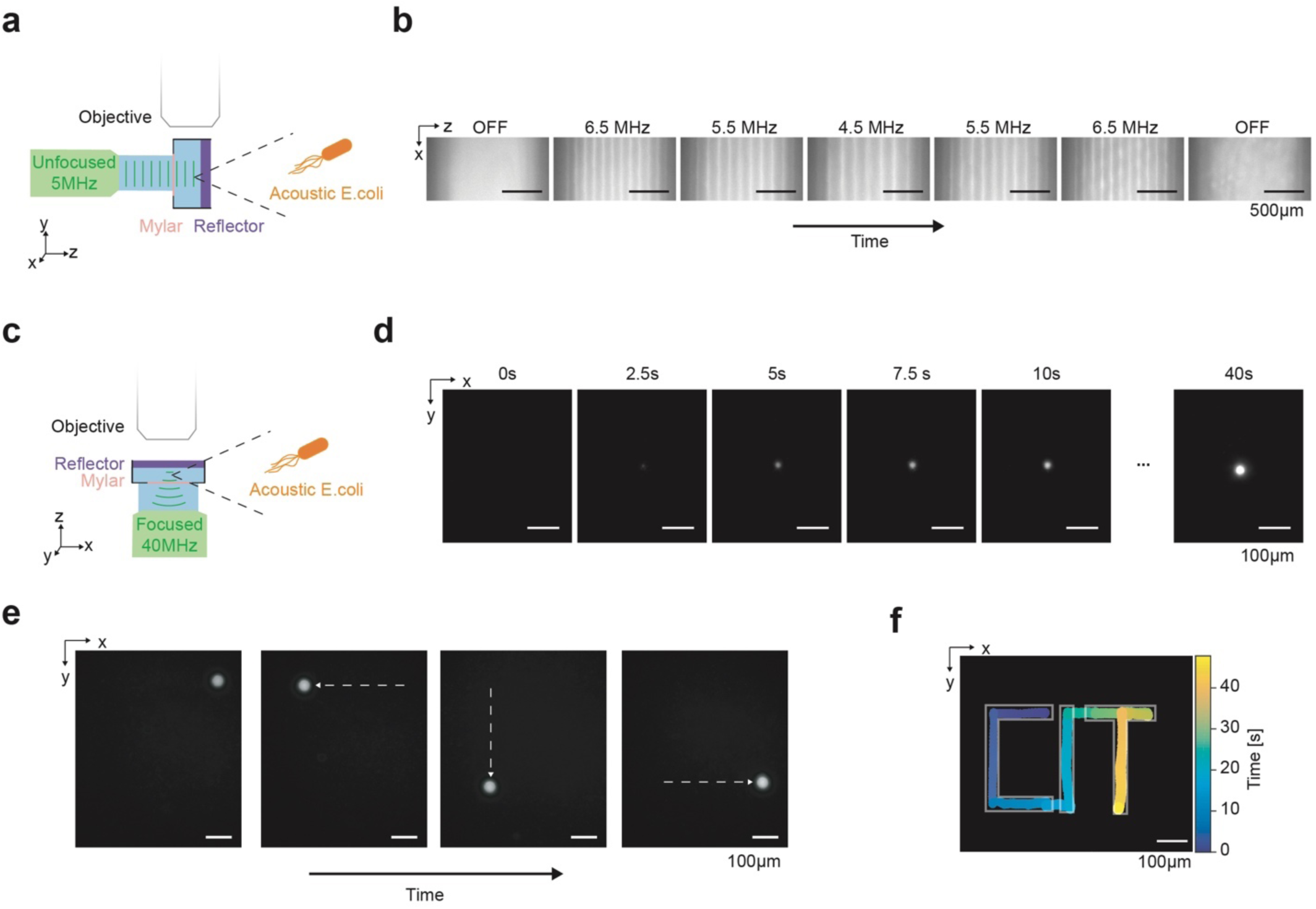
Dynamic patterning of acoustic bacteria. **a**, Diagram of the acoustic chamber setup for frequency-controlled spatial patterning. A transducer is aligned orthogonal to a glass reflector using a 3D-printed holder. The sound wave passes through a mylar membrane, is reflected by the glass reflector, and forms a standing wave near the reflector. The sample region containing acoustic *E.coli* is imaged using an epifluorescence microscope. **b**, Sequential fluorescence images of acoustic *E.coli* in the presence of an acoustic standing wave at varying frequencies. Frequencies were changed every 50 seconds. **c**, Diagram of the acoustic chamber setup for image-guided trapping and positioning of acoustic *E.coli*. Imaging is performed along the axis of a focused 40 MHz transducer. **d**, Sequential fluorescence images of the formation of a cluster of acoustic *E.coli* at the ultrasound focus. **e**, Fluorescence images of a cluster of acoustic *E.coli* positioned at distinct locations in the x-y plane. The positioning is controlled by the translation of the transducer in the x-y plane using a micromanipulator and is guided by real-time fluorescence imaging of the bacteria. **f**, Overlaid positions of the cell cluster, color-coded by time, to form a spatiotemporal pattern writing out “CIT”.

Another method of acoustic manipulation involves the confinement of acoustic particles at the focus of an ultrasound transducer^26–29^, allowing the particles to be concentrated and transported between discrete locations in space, analogous to an optical trap. To determine whether focal trapping is possible with engineered acoustic cells, we generated a trap using a 40 MHz focused ultrasound transducer reflected on glass (Fig. 5c). This configuration is expected to exert radial ARF on the cells towards the center of the ultrasound focus. As expected, GV-expressing cells within this acoustic field coalesced into a cellular cluster upon ultrasound application (Fig. 5d and **Supplementary Movie 4**) and could then be moved around in space by laterally translating the ultrasound transducer, generating a desired spatiotemporal pattern (Fig. 5, e-f and **Supplementary Movie 5**). These results demonstrate the ability of genetically encoded GVs to specifically sensitize engineered living cells to acoustic separation, trapping, patterning and dynamic rearrangement.

## DISCUSSION

Our results establish GVs as the first biomolecules to be directly manipulated and patterned with ultrasound. Due to their unique physical properties, GVs produce the highest acoustic contrast of any stable particle in an aqueous environment, allowing these nanostructures to experience strong ARF despite their sub-micron size. Furthermore, the expression of GVs inside engineered cells greatly enhances and changes the sign of the force experienced by these cells due to ultrasound, enabling the selective acoustic manipulation and patterning of these cells based on their genotype.

This technology is expected to find applications in several areas of biotechnology and biomaterials. First, the ability of GVs and GV-expressing cells to be patterned and manipulated dynamically in 3-D space will enable the development of protein- and cell-based materials for applications in tissue engineering^1^ and living materials^21,22^. In these applications, ultrasound has intrinsic advantages compared to optical, magnetic or printing-based approaches due to its compatibility with opaque media, fine spatial resolution and non-invasive access. Second, the development of acoustofluidic^6,30^ devices combining ultrasound with microfluidic channels creates opportunities for GVs to be used to as nanoscale acoustic labels, binding to and driving the separation of specific components of complex biological samples. To this end, GVs are readily functionalized with moieties providing the ability to bind specific biomolecular targets^8,13^. Likewise, the ability of GVs to serve as genetically encoded acoustic enhancers will allow their expression to designate specific cells for separation, trapping and patterning using ultrasound. These capabilities could be extended from *in vitro* devices to inside living animals or patients using emerging approaches for *in vivo* ARF^31^. Finally, GVs could be used as a nanoscale actuator to locally apply specific forces to biological systems, which may be useful for studies of endogenous mechanosensation or for engineered mechanisms of non-invasive cellular control^5^.

While our findings have expected utility in a broad range of contexts, additional studies are needed to fully characterize and further expand the capabilities of GVs as transducers of ARF. First, it will be useful to build on the fundamental demonstrations in this study by applying GVs to specific biological problems, taking advantage of their potential for biomolecular and genetic engineering. Second, the theoretical model of GV acoustic contrast used in this study approximated that GVs have spherical geometry and that their shell has a constant density and compressibility as a function of applied acoustic pressure. In reality, GVs are anisotropic cylindrical nanostructures that can undergo reversible buckling under applied acoustic pressure^32,33^. This buckling behavior is expected to enhance the effective compressibility of GVs and thereby the ARF they experience. Theoretical analysis of GV ARF with more realistic geometry and experiments using a broader range of pressures encompassing the buckling regime could inform the engineering and use of these biomolecules in ARF applications. Third, it will be useful to explore the inter-particle interactions arising between GVs and GV-expressing cells in an applied acoustic field, as this may influence their clustering, separation and motion. Based on these additional physical insights, it may be possible to genetically engineer new GV phenotypes with size, shape and mechanical properties enhancing their exceptional response to ARF and further propelling the fantastic voyage of engineered biomolecules and cells in biomedicine and biomaterials.

## MATERIALS AND METHODS

### Estimation of acoustic contrast factor

Acoustic contrast factors were calculated using the equation:

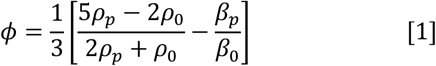

where ρ_*p*_ and ρ_0_ are the density of the particle and the fluid, respectively, β_*p*_ and β_0_ the compressibility of the particle and the fluid, respectively. Values of ρ_*p*_ and β_*p*_ for GVs were obtained from literature ^14,15^. Values of ρ_*p*_ and β_*p*_ for the acoustic *E.coli* were obtained by assuming that 10% of the intracellular space was occupied by GVs^12^ and calculating the volume-averaged density and compressibility.

### Preparation of gas vesicles

GVs from *Anabaena flos-aquae* (Ana), *Bacillus megaterium* (Mega), and Ana GVs with GvpC removed (AnaΔC) were prepared as previously described.^34^ Dylight415-Co1 N-hydroxysuccinimide ester (Thermo Fisher Scientific) was reacted with GVs in PBS for 2 hours at 10,000:1 molar ratio, protected from light, on a rotating rack. 10 mM Tris buffer was then added to the solution to quench unreacted dye. Labeled GVs were subjected to dialysis and buoyancy purification. Pre-collapsed GVs controls were prepared by application of hydrostatic pressure in a capped syringe.

### Preparation of acoustic E.coli

GV-expressing cells were produced by transforming a pET28a plasmid containing the *arg1* gene cluster^12^ (Addgene #106473) into BL21(A1) *E. coli* (Thermo Fisher Scientific). The transformed cells were first grown overnight at 37 °C in LB media supplemented with 1% glucose, and subsequently diluted 1:100 into LB media supplemented with 0.2% glucose. When the optical density at 600 nm (OD600) of the culture reached between 0.4 and 0.6, 400 μM IPTG and 0.5% l-arabinose were added to induce the expression of GVs. The expression proceeded at 30 °C for 22 hours. High-expressing cells were enriched by centrifugation-assisted floatation at 300 g. Cell density was measured after collapsing any intracellular GVs to eliminate their contribution to optical scattering. *E.coli* with pre-collapsed GVs were prepared by application of hydrostatic pressure to the cell culture in a capped syringe. Fluorescently labeled bacteria were prepared by incubating the cells with 10 µM of Baclight Green bacterial stain (Thermo Fisher Scientific) for 40 minutes at room temperature, protected from light, and followed by two rounds of buoyancy purification to remove excess dye.

### Acoustofluidic setup

The acoustofluidic channel was designed in SolidWorks, and fabricated in a clean room facility following a protocol modified from one previously described^35^. Briefly, AZ1518 positive photoresist (Merck) was patterned onto a <100> silicon wafer (University Wafer) using a photomask, and developed in AZ340 solution. 50 cycles of deep-reactive ion etching (PlasmaTherm, SLR Series) were used to etch the channels into the wafer. The channel depth was measured using a profilometer (P15, KLA-Tencor). The photoresist was then removed, and the wafer was cleaned with piranha solution. A Borofloat 33 borosilicate glass wafer was anodically bonded to the silicon overnight at 500V, 400°C using a custom setup. Inlet holes were drilled through the glass layer using a diamond drill bit (Drilax) and joined with microfluidic connectors (Idex Health & Science) using Epoxy (Gorilla). A custom PZT-5A piezoelectric element (American Piezo Company) was attached to the silicon beneath the channel using cyanoacrylate (Loctite). The input signal to the PZT was programmed in MATLAB and generated using an arbitrary waveform generator (Tabor Electronics). The output waveform was validated by an oscilloscope (Keysight Technologies) before being amplified by an RF power amplifier (Amplifier Research) and connected to the PZT. The samples inside the channel were imaged using a custom-built upright epifluorescence microscope with an LED source (Thorlabs) and a sCMOS camera (Zyla 5.5, Andor).

### Single-particle tracking experiment and analysis

Fluorescently labeled GVs, suspended in buffer (DI water, 0.01% v/v Tween-20), were introduced into the acoustofluidic channel via a syringe. The background flow was naturally slowed until particles stayed within the field of view longer than the acquisition time of approximately 2 minutes. The particles were then imaged at 20 frames per second for approximately 20 seconds before ultrasound was turned on. The ultrasound was then turned on (3.75 ± 0.1MHz sweep, 1 ms sweep repetition time, 3.8V peak-to-peak, continuous wave) for approximately 100 seconds. Pressure-collapsed GVs, and 200 nm diameter fluorescent polystyrene particles (Thermo Fisher Scientific) were subjected to the same procedure.

Particle detection was performed in ImageJ using the MOSAIC ParticleTracker plugin to obtain time-dependent particle coordinates in the direction towards the walls, *x*(*t*). Particle trajectories were exported and analyzed in MATLAB using custom scripts. The coordinates were split into before-ultrasound and during-ultrasound groups. Only particles with trajectories in both groups were included in the analysis.

Trajectories during the Brownian period were used to calculate the mean-squared-displacement, < ∆*x* >^2^, for different time durations, ∆*t*. Linear regression was used to extract the diffusion coefficient, *D*, for each particle following the relationship < ∆*x* >^2^= 2*D*∆*t*. The mobility, μ, of the particle was then obtained using the Einstein relation:

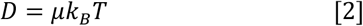

where *k*_*B*_ is the Boltzmann constant, and *T* the temperature. Trajectories recorded during the ultrasound period were fitted to an equation of motion accounting for the sinusoidal pressure profile to obtain the peak particle velocity in the acoustic field. Given the profile of the pressure in the channel *P* (*x*, *t*) = *P*_*peak*_ cos(*kx*) sin(ω*t*), where *k* is the wave number and ω the angular frequency, the radiation force, *F*_*ARF*_, acting on the particles is:

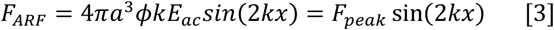

here *a* is the particle radius, ϕ the acoustic contrast factor, 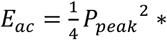 *β*_0_ the acoustic energy density, and *F*_*peak*_ the peak ARF^16^.

At low Reynold’s number, *F*_*ARF*_ = *F*_*drag*_ ∝ *v*_*p*_, where *F*_*drag*_ is the drag force and *v*_*p*_ the particle velocity. Therefore, *v*_*p*_ = *v*_*peak*_ sin(2*kx*), where *v*_*peak*_ is the peak particle velocity. The particle position, *x*_*p*_(*t*), over time within an acoustic field is thus related to the peak velocity by:

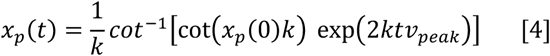

Fitting the particle trajectory to this equation allowed us to obtain *v*_*peak*_. Combining the particle mobility μ and the peak velocity *v*_*peak*_, the peak ARF was calculated using 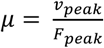. The hydrodynamic radius *a*_*H*_ of the particles was determined using the Stokes-Einstein equation:

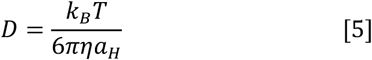

where η is the solution viscosity. Fitting the force measurements to a fractal clustering model

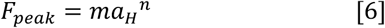

to obtain the scaling coefficient m, and the force-mobility exponent n, the peak ARF for a single GV, *F*_*peak_sGV*_, was calculated by substituting the average hydrodynamic radius of a GV^34^, *a*_*H_sGV*_ = 125 *nm*. The acoustic contrast factor of a single GV, ϕ_*sGV*_, was then obtained using the equation:

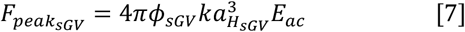

where *E*_*ac*_ is the acoustic energy density of the applied ultrasound, as determined by a separate calibration. Finally, this equation is used to predict the peak ARF for a single GV at various acoustic parameters.

### Acoustic GV collapse in microfluidic channel

A syringe pump was used to introduce fluorescently labeled AnaΔC GVs into the acoustofluidic chip at a controlled flow rate of 0.5 µl/min. Fluorescence images were acquired while the PZT was driven at three different voltages. The acoustic energy density for the three trials was kept constant by choosing the appropriate duty cycle according to *Duty Cycle* * *Voltage*^2^ = *constant*. A steady-state image was selected and was projected onto the x-axis to determine the locations of the discontinuity in the fluorescence signal. The location was marked with the critical collapse pressure of AnaΔC of 0.6 MPa, and the acoustic pressure in the entire channel was calculated by assuming a sinusoidal pressure profile with antinodes at each wall.

Fluorescently labeled Mega GVs were introduced into the channel in a similar manner and subjected an acoustic field with a peak acoustic pressure of 1.2 MPa, as measured using the collapse profile of AnaΔC.

### Acoustic manipulation of bacteria in microfluidic channel

Fluorescently labeled *arg1*-expressing *E. coli* and pre-collapsed controls, prepared as described above, were suspended in PBS and loaded into the acoustofluidic channel described above. Continuous wave ultrasound was applied at 3.75 MHz, 7.6 V peak-to-peak. Images of the channel were acquired for 10 seconds during ultrasound application as described above.

### Dynamic patterning of acoustic bacteria

An acoustic setup was built to generate a standing wave with reconfigurable wavelengths, by reflecting the sound generated by a single-element transducer (V310, Olympus) off a glass coverslip (VWR). A holder was designed in SolidWorks and 3D-printed (3D Systems) to facilitate the alignment of the transducer with the reflector and to create a sample chamber sandwiched between the reflector and an acoustically transparent mylar membrane (Chemplex, 2.5 μM thickness). The acoustic setup was placed into a water bath to provide acoustic coupling between the transducer and the sample chamber, and fluorescently labeled *arg1*-expressing *E. coli* prepared as above were suspended in PBS and loaded into the sample chamber. Ultrasound (continuous wave) was applied to the sample, and fluorescent images were acquired with the imaging plane parallel to the sound propagation axis. The ultrasound frequency was varied between 4.5 and 6.5 MHz in 1 MHz steps every 50 seconds.

### Image-guided positioning of acoustic bacteria

For radial acoustic trapping and movement, a sample dish was created allowing the placement of the image plane orthogonal to the sound propagation axis. The glass bottom of a 35-mm glass-bottom petri dish (Matsunami) was removed using a glass cutter and replaced with a Mylar film. *arg1*-expressing *E. coli* prepared as above and suspended in PBS were added to the center of the dish, and sealed using a glass coverslip. A 40 MHz focused single-element transducer (V390-SU/RM, Olympus) was mounted onto a micromanipulator and positioned beneath the dish. To align the transducer with the glass reflector, the transducer first emitted 5-cycle pulses and received the echo from the glass coverslip. The amplitude of this echo was maximized by adjusting the position of the transducer using the micromanipulator. To trap the acoustic bacteria, the transducer was then driven with a continuous wave 40 MHz input while fluorescent images were acquired. After a cell cluster was formed in the center of the acoustic focus (**Supplementary Figure 4**), the transducer was moved in the x-y plane using the micromanipulator, guided by the optical image, to form the desired positioning sequence.

### Statistical analysis

Statistical methods are described in each applicable figure caption. Measured values are stated in the text as the mean ± the standard error of the mean. Standard error propagation methods were used where appropriate.

## ACKNOWLEDGEMENTS

The authors thank James Friend and Aditya Vasan for helpful discussion, Hunter Davis for help with fluorescence microscopy, Zhiyang Jin for assistance with bacteria protein expression, Gabrielle Ho for assistance with electron microscopy, and Xiaozhe Ding for contribution to initial experiments. This work was funded by the National Institutes of Health (R01EB018975 to MGS), the Pew Scholarship in the Biomedical Sciences (to MGS) and the Packard Fellowship in Science and Engineering (to MGS). DW was supported by a Medical Engineering Amgen Fellowship. DB was supported by Newton International Fellowships (NF 161508). DMaresca was supported by the Human Frontiers Science Program Cross-Disciplinary Fellowship. MPA was supported by A*STAR.

## AUTHOR CONTRIBUTIONS

DW, MGS, DB, and DMaresca conceived the study. DW and DB designed, planned, and conducted the experiments. DW, CC, and DRM designed and fabricated the acoustic devices. DW and DMalounda prepared the samples. DMaresca and MPA contributed to experiments. DW analyzed the data. DW and MGS wrote the manuscript with input from all authors. MGS supervised the research.

## COMPETING INTERESTS

The authors declare no competing financial interests.

