## Supplementary Information for "Genetically encoded nanostructures enable acoustic manipulation of engineered cells"

### Supplementary Figures

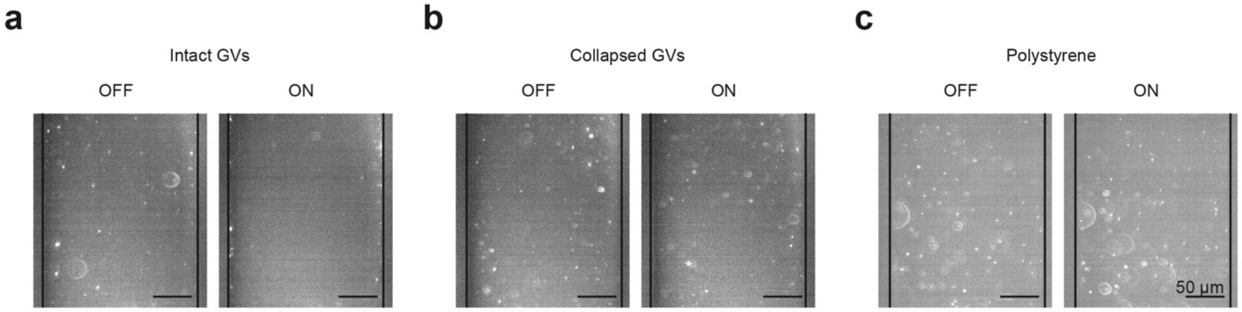


**Supplementary Fig. 1 | Control particles do not experience substantial ARF.** Fluorescence images of intact GVs (**a**), pressure-collapsed GVs (**b**), and polystyrene nanoparticles (**c**) inside the microfluidic channel before ultrasound (OFF) and 100 seconds after ultrasound has been turned on (ON). Device and acoustic conditions are as described in Fig. 2.

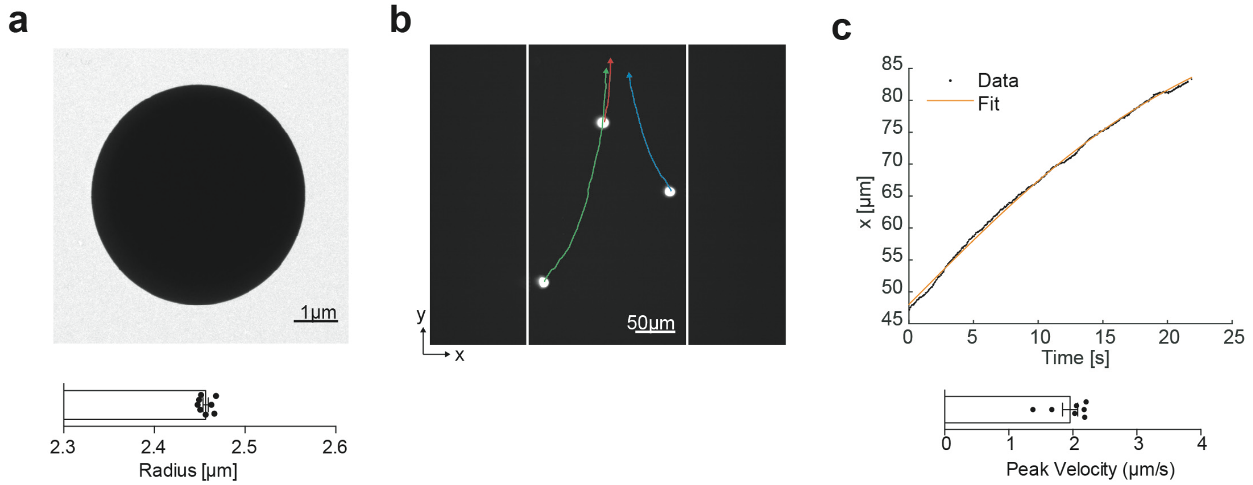


**Supplementary Fig. 2 | Calibration of the acoustic energy inside the acoustofluidic channel.** **a**, Representative TEM image of a polystyrene particle (top) and quantification of the particle radius (bottom, 2.457±0.003 μm, mean±S.E.M., n=7). **b**, Fluorescence image and overlaid acoustophoretic trajectory of polystyrene particles inside the acoustofluidic channel. The white lines demarcate the edges of the channel. Arrows indicated direction of particle movement. **c**, Representative single-particle trajectory in the x-direction during ultrasound stimulation (top), and quantification of the peak particle velocity (bottom, 2.0±0.1 μm/s, mean±S.E.M., n=7). The acoustic energy is determined using the radius, peak velocity, and the acoustic contrast factor of polystyrene particles (**Fig.1e**).

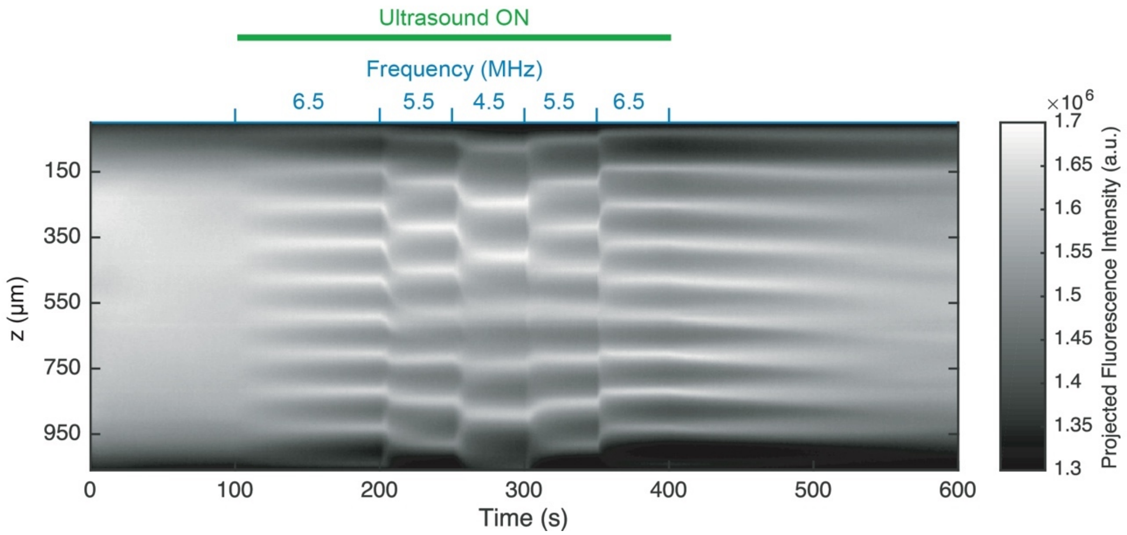


**Supplementary Fig. 3 | Cell patterns can be reconfigured on the timescale of seconds.** Kymograph of projected fluorescence signal from *arg1*-expressing *E.coli* during the application of ultrasound at different ultrasound frequencies. Conditions as described in **Fig. 5, a-b**.

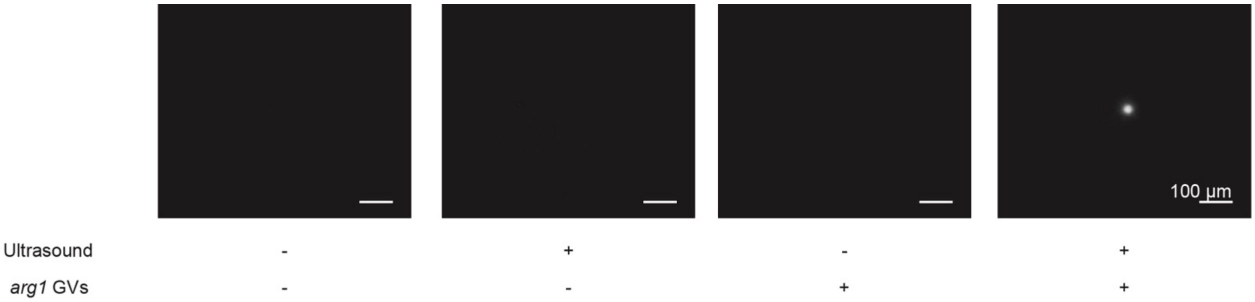


**Supplementary Fig. 4 | Bacteria cluster formation requires intact intracellular GVs.** Fluorescence images of arg1-expressing E.coli with intact (+) and collapsed (-) intracellular GVs before and 40 seconds after ultrasound application.
